# Ecotype-specific genomic features within the *Escherichia* cryptic clade IV

**DOI:** 10.1101/2024.07.17.603965

**Authors:** Martín Saraceno, Nicolás Frankel, Martín Graziano

## Abstract

*Escherichia* cryptic clades represent a relatively unexplored taxonomic cluster believed to exhibit characteristics associated with a free-living lifestyle, which is known as the environmental hypothesis. This hypothesis suggests that certain *Escherichia* strains harbour traits that favour their environmental persistence, thus expanding the ecological commensal niche of the genus. While surveying *Escherichia* diversity in an urban South American stream we isolated the first environmental cryptic clade IV strain in South America (339_SF). Here we report the genomic characterization of 339_SF strain in the context of existing genomic information for cryptic clade IV. A comparative analysis of genomes within the same clade stemming from diverse ecological sources and geographical locations reveals close phylogenetic proximity between our isolate and strains of environmental origin. In the genomes of cryptic clade IV strains that were isolated from environmental niches we observed enrichment of functional genes related to responses to adverse environmental conditions and a low number of genes with clinical relevance among. Our findings highlight substantial intra-group genomic structuring linked to ecological origin and shed light on the genomic mechanisms underlying the naturalization phenomena within the *Escherichia* genus.

## Introduction

The *Escherichia* genus includes widely known species such as *E. coli, E. fergusonii* and *E. albertii*, and five monophyletic groups named “cryptic” clades I to V (Walk, 2015). The name of the latter is based on the inability to distinguish them from representative isolates of *E. coli* through typical biochemical diagnostic reactions. However, relatively recent phylogenomic analyses have shown that these lineages are divergent from stem members of the genus (Walk *et al*., 2009; Luo *et al*., 2011). Previous studies associated some cryptic clades with a free-living lifestyle, a phenomenon known as the *environmental hypothesis*. The initial evidence for this proposition was based on the overrepresentation of these clades in environmental samples (Walk *et al*., 2007, 2009). Subsequent phenotypic, genomic and transcriptomic analyses supported the link as well (Walk, 2015; Di Sante *et al*., 2018), but the evidence behind this hypothesis is still limited.

Cryptic clade I is closely related to *E. coli* and genomic studies indicate that both groups carry common virulence factors, which suggests that clade I strains could also be as pathogenic as some *E. coli* strains (Steinsland *et al*., 2010; Walk, 2015). Bacterial virulence factors encode genes that facilitate infection, such as toxins or proteins needed for bacterial adherence (Holden *et al*., 2009; Stecher and Hardt, 2011; Acosta-Dibarrat *et al*., 2021). The presence of virulence and antibiotic resistance factors in bacterial genomes is often linked to specific selection pressures associated with hosts (Jernberg *et al*., 2010; Becattini *et al*., 2016). On the other hand, field, experimental and genomic evidence support the hypothesis that the remaining clades (II to V) survive outside animal hosts (Di Sante *et al*., 2018). This assumption is based not only on the fact that they have been generally isolated from soil or surface water (they have not been linked to cases of human or animal infection), but also on the lack of virulence factors for intestinal and extra-intestinal infection (Ingle *et al*., 2011; Vignaroli *et al*., 2015). Moreover, a comparative genomic study suggested that genomes of cryptic clades associated with a free-living style are enriched in genes that confer greater fitness in the environment, e.g. contributing to novel metabolic pathways for the exploitation of alternative energy sources (Luo *et al*., 2011). A caveat of the analyses mentioned above is that they were performed with only a few genomes. Thus, a more comprehensive analysis that includes all currently available genomes is in need for a better understanding of the genomic signatures related to a free-living style.

Addressing the ecotype-genetic relationship within the genus *Escherichia* is relevant due to the increasing number of studies that have reported the ability of certain *E. coli* strains to persist in secondary habitats (Ishii *et al*., 2006; Lee *et al*., 2006; Mackowiak *et al*., 2018). Thus, the review of the traditional niche assigned to *E. coli* is framed by a larger discussion on the ecological history of the genus itself, which calls for an increase in genomic and phenotypic characterizations. It has been hypothesized that the phenomenon of environmental persistence could be the result of a complex balance between the differential fitness of some *E. coli* strains and the occurrence of permissive ecological conditions, i.e., optimal physicochemical characteristics, nutrient availability, low competition, among others (Surbeck *et al*., 2010; Jang *et al*., 2017). Furthermore, secondary habitats such as urban streams, where point and diffuse sources of bacterial input usually coexist with the native microbiome and temperatures are usually benign and nutrient availability high, are hot spots for genetic exchange and the emergence of new strains, which can be pathogenic (McLellan *et al*., 2015). In this line, previous research has detected the imprint of isolation sources both in genomic traits of epidemiological importance, as well as in phenotypes linked to long-term persistence in *E. coli* (Berthe *et al*., 2013). In a recent study, also on *E. coli*, associations among genetic backgrounds and specific habitats were uncovered and horizontal gene flow was found to be an important mechanism driving the reinforcement of gene structuring (Touchon *et al*., 2020). Due to the sanitary and ecological interest of these phenomena, we consider that it is relevant to explore the genomic features associated with a free-living style within the genus *Escherichia*.

In a previous work, we reported the isolation of a member of cryptic clade IV from a South American urban stream (Saraceno *et al*., 2020). Here we report the genomic characterization of this strain, while performing a comparative analysis of genomes within the clade IV from diverse ecological and geographic sources. We present evidence of intra-clade IV functional genes structuring linked to the ecotype of origin. We discuss our results within the framework of the environmental hypothesis and the occurrence of niche-specific selective pressures.

## Methods and Materials

### Environmental cryptic clade IV isolation and phylogenetic assignment

The isolate 339_SF, belonging to the cryptic clade IV, was isolated, cryopreserved and phylogenetically characterized as previously mentioned in Saraceno *et al*. (2020). Briefly, environmental isolates were obtained from the water column of San Francisco urban stream (Buenos Aires, Argentina) by culturing methods: a first-round employing Chromocult® Coliform Agar selective medium (MilliporeSigma) and, in a second step, blue- to purple-coloured selected colonies were streaked at least two times onto Levine E.M.B. (Eosin Methylene Blue) agar plates to assure isolates purity. The phylogenetic assignment was carried out through a series of multiplex PCR procedures. A first round, through the amplification of the *araA, chuA, yjaA* and *TspE4*.*C2* genes, which determined that isolate 339_SF belonged to an *Escherichia* cryptic clade, and a subsequent second round of a double PCR method, based on the amplification of *aes* and *chuA* genes, to finally determined its cryptic lineage membership (Clermont *et al*., 2011, 2013). All PCR procedures were carried out using a loaded loop after ringing on each colony as a template, in a 20 μL final volume, and 10 μL of 2X GoTaq® Green Master Mix (Promega), following respective literature indications.

### DNA extraction and sequencing

Before DNA extraction, 339_SF isolate was cultured overnight in Luria Bertani broth at 37°C and 1,5 ml of this culture was pelleted by ultracentrifugation (13000 rpm for 5 min). Obtained pellet was incubated with saline EDTA buffer (C_10_H_16_N_2_O_8_ 0.01 M and NaCl 0.15 M; pH = 8.0) and proteinase K. DNA purification was held through serial washes with chloroform:isoamyl alcohol (24:1), precipitated with isopropyl alcohol and re-suspended in low-TE (Tris 1 mM and EDTA 0.1 mM; pH = 7.0). Genomic DNA integrity was assessed through agarose gel electrophoresis (1% agarose). DNA mass was estimated by the inclusion of a mass ladder (MassRulerTM DNA Ladder Mix) and its concentration by a Qubit assay (dsDNA Quantitation, broad range). Whole genome sequencing was performed in Novaseq platform (Illumina, 2×150 paired-end), obtaining a Q30 of 91.99% and an estimated sequencing coverage of 890X.

### Genome assembly and gene annotation

Reads quality was assessed using FASTQC and Kraken2 (de Sena Brandine and Smith, 2019; Wood *et al*., 2019). Genome assembly was made using Unicycler (Wick *et al*., 2017). The quality of the assembly was assessed by Quast (Gurevich *et al*., 2013) and its phylotyping was corroborated using Clermontyping *in silico* method (Beghain *et al*., 2018). N50 was of 536699 bp and L50 of 3 contigs, revealing a successful assembly procedure. Gene annotation was done using Prokka (minimum contig size 300 bp, genus-specific BLAST databaseforg. *Escherichia*) (Seemann, 2014). Search for genome features of clinical and epidemiological interest was done using ABRicate tool and VFDB database for virulence factors, NCBI and Resfinder databases for antimicrobial resistance (AMR) genes and Plasmidfinder database for plasmids replicons (Carattoli *et al*., 2014; Seemann, 2020; Florensa *et al*., 2022; Liu *et al*., 2022), considering only those with both coverage and identity above 95%.

### Phylogenetics

Genome assemblies were obtained from public and curated databases such as EnteroBase (Zhou *et al*., 2020), belonging to the cryptic clade IV (44 genomes) and the cryptic clades I (2 genomes), II (1 genome), III (2 genomes) and V (2 genomes). Also, genome assemblies from *E. albertii, E. fergusonni, E. coli* K-12 and *E. coli* main pathogenic groups were acquired: EPEC (enteropathogenic *E. coli*), ETEC (enterotoxigenic *E. coli*), EIEC (enteroinvasive *E. coli*), STEC (Shiga toxin-producing *E. coli*), EAEC (enteroaggregative *E. coli*), EHEC (enterohemorrhagic *E. coli*), UPEC (uropathogenic *E. coli*) y NMEC (neonatal meningitis-causing *E. coli*). Among clade IV genomes, 10 belong to the environmental isolation ecotype, 14 to human hosts, and 20 to other hosts (wild and domestic animals); all of them from different geographic locations around the world. Accession information and metadata from the 62 genomes are listed in **Table S1**. All the assemblies were analyzed following the same paths of annotation and search for genomic features of clinical interest mentioned above. From the annotation files obtained, a pangenome analysis was conducted through Roary pipeline (Page *et al*., 2015), identifying the core and accessory genes among genomes. Core genes were employed as input for the construction of the phylogenetic relationships. The maximum likelihood method with a bootstrapping of 1000 was employed for the inference of the tree using IQ-TREE (Nguyen *et al*., 2015). Lastly, the resulting phylogenetic tree was rooted in the midpoint and the graphics were edited using FigTree and Inkscape, respectively (Harrington, 2004; Rambaut, 2018).

### Pangenome of cryptic clade IV and genetic enrichment analysis according to ecotype

The set of annotated genomes belonging to the cryptic clade IV (44) was used as input to design a presence/absence matrix of functional genes - 3524 entries-, as characterized by UniProtKB database (The Consortium Uniprot, 2023). Genes detected in only one genome or across the entire genome set were discarded, and a hierarchical clustering analysis was held with the resulting matrix. Further, the set was subdivided into two subsets according to its ecotype host/environmental origin. The number of times each gene was detected in each group was counted (repeated copies of the same gene per genome were considered only one time). Consequently, the detection frequencies of each gene were calculated inside each subset. To consider a gene as enriched within one of the two ecotypes, a mixed criterion was used based on the resulting frequencies: a minimum threshold of 80% detection in one of the subsets was required and, jointly, a detection frequency of less than 80% in the complementary subset. After the identification of the genes considered enriched in each ecotype, a review of the literature was carried out to assign them a functional and ecological context.

## Results

### Genomic features of the environmental cryptic clade IV isolate

Strain 339_SF was isolated from an urban stream in South America and characterized as a member of the cryptic clade IV (Saraceno *et al*., 2020). To our knowledge, there is no previous report of an environmental isolate belonging to *Escherichia* cryptic clades in the region. We sequenced the whole genome of strain 339_SF and produced a total of 29 contigs with 4283 predicted coding sequences. The total length of the assembly (4637586 bp) and the percentage of GC content (50.86%) are consistent with the expectations for an *Escherichia* genome (Hildebrand *et al*., 2010). No genes associated with antibiotic resistance or plasmid replicons were detected, while a total of 16 sequences matched virulence factors (VFs). Among the VFs detected, none are strictly associated with a defined *E. coli* pathotype. For example, we found factor *sat*, which is often associated with UPEC and EAEC profiles, but which is distributed between commensal strains as well (Toloza *et al*., 2015). In addition, we detected the numerous variants of the type 1 fimbria precursor gene (*fimB, fimC, fimD, fimG* and *fimI*), which are common VFs in strains associated with UPEC infections (Müller *et al*., 2009). We also identified factors associated with siderophore function (*chuX,fepA, fepB, fepD*and *entS*), which are widespread among various *E. coli* strains and with presumed negative effects on hosts (Ozenberger *et al*., 1987; Bleuel *et al*., 2005; Suits *et al*., 2009) and factors that encode components of the general gram-negative secretion pathway (*gspG, gspH, gspI* and *gspL*) (Py *et al*., 2001). *KpsD*, a key factor for the synthesis of the capsule in *E. coli*, which is the structure that confers resistance during extraintestinal infections (Russo *et al*., 1998; Duan *et al*., 2020), was also detected.

### Genes of clinical interest in Clade IV genomes

We compared available genomes of cryptic clade IV with that of 339_SF. Among the 44 genomes, the minimum number of VFs detected was 10 and the maximum was 23, with an average of 14.8. Of the 43 VFs found across the set, 12 were shared by 84.4% of the clade IV genomes. These 12 factors are present in the genome of 339_SF (see **Table S2** for the genomic traits of clinical and epidemiological relevance). The genomes carrying more VFs were isolated from humans from different continents (ESC_BA8399AA, ESC_EA6501AA and ESC_LA2398AA, with 23 putative genes each, respectively from Africa, Europe and South America). We did not detect AMR genes in the 339_SF genome, as in most of the cryptic clade IV genomes (39 of 45 genomes have no resistance determinants). However, we detected 13 factors in the genome of a poultry isolate (ESC_FA7484AA). Similar results are observed when looking at plasmid replicons: clade IV genomes carry one plasmid replicon on average, and almost half of them do not carry replicons. Thus, 339_SF possesses a similar profile of factors of clinical interest when compared to other cryptic clade IV genomes from various geographic or ecotype origins.

### Phylogenomics of Cryptic Clade IV

We performed a phylogenetic analysis based on *core* genes with the available genomes of clade IV, genomes of reference strains from the rest of the cryptic clades and other main taxa of the genus *Escherichia*. After annotating genes across all assemblies, 501 were found in the entire set (**Table 1**). Clade IV strains formed a monophyletic group in the tree (**Figure 1**). Genomes belonging to clades II, III and V appear close to clade IV. Clade I genomes fall close to *E. coli* strains (K-12 and those considered pathogenic) and *E. fergusonii*. 339_SF groups with a subset of clade IV genomes that include 6 other isolates of environmental origin. Furthermore, this cluster of genomes tends to lack antibiotic resistance genes and plasmid replicons, while carrying a similar number of VFs - between 14 and 16- (**Table S2**).

**Table 1.** Abundance of gene types according to their sharing percentage between the strains.

| Types of genes | Percentage sharing between strains | Number of genes |
| --- | --- | --- |
| *core* | 99% <= genomes <= 100% | **501** |
| *soft core* | 95% <= genomes <= 99% | 1604 |
| *shell* | 15% <= genomes <= 95% | 2493 |
| *cloud* | 0% <= genomes <= 15% | 18485 |
| Total | 0% <= genomes <= 100% | **23083** |

**Figure 1:**
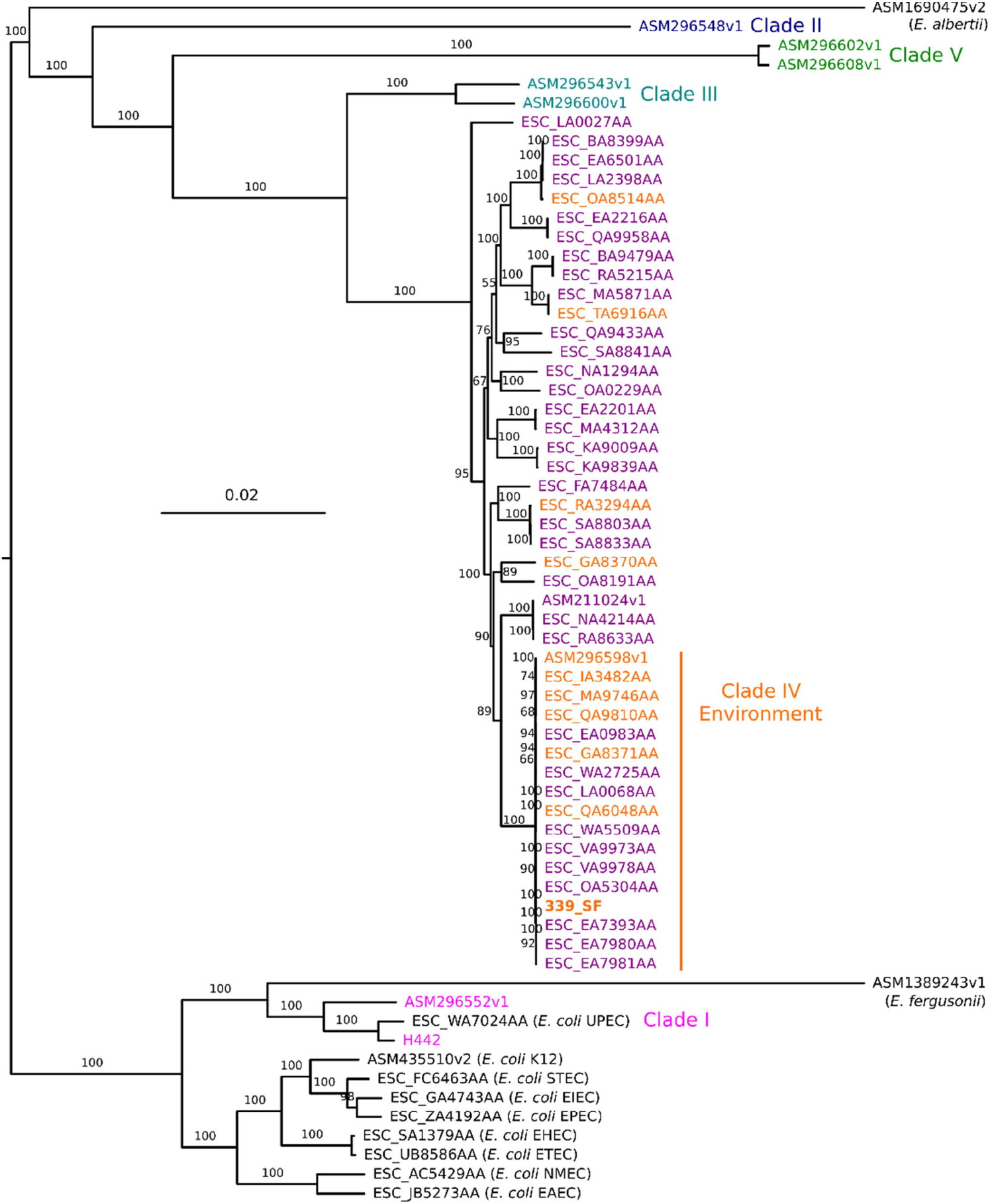
Phylogenetic relationships amongst selected *Escherichia* genus strains. Genomes from cryptic clades IV (ecotypes are discriminated by colour, host -orange- or environmental -violet-), I, II, III and V, E. coli K12 and reference strains from its main pathotypes, as well as *E. fergusonii* and *E. albertii* were included. 339_SF is labeled in bold orange and the cluster containing relatively more environmental ecotype genomes is indicated. Full sequences from the 501 core genes were aligned presenting a base overlap region of 0.4 Mb and phylogeny was conducted as detailed in Materials and Methods. Bootstrap support values are indicated on each branch.

### Association between genomic features and ecotype within cryptic clade IV

We grouped genomes following a functional genetic content perspective to further investigate the relationship between genomic features and ecotype within cryptic clade IV. Hence, a gene presence/absence matrix was created using a pangenome (the totality of annotated genes with experimentally validated or inferred functionality across all the set of clade IV genomes). After purging the matrix of genes detected in a single genome and present in all genomes, we retained 870 entries. A hierarchical clustering based on Pearson’s positive correlations was performed with these data (**Figure 2**). The tree shows that CC IV genomes are grouped into two main clusters of 18 and 26 genomes. Globally, this organization is similar to that observed in the phylogeny based on core genes (**Figure 1**). 339_SF is included in the group that contains the highest proportion of isolates of the environmental ecotype (7 of 18 genomes, versus 4 of 26). We also observed that 13 of the genomes within the cluster of 18 have a very strong positive correlation (upper right corner, **Figure 2**). 339_SF and 4 other genomes from environmental origin are part of these highly correlated genomes.

**Figure 2:**
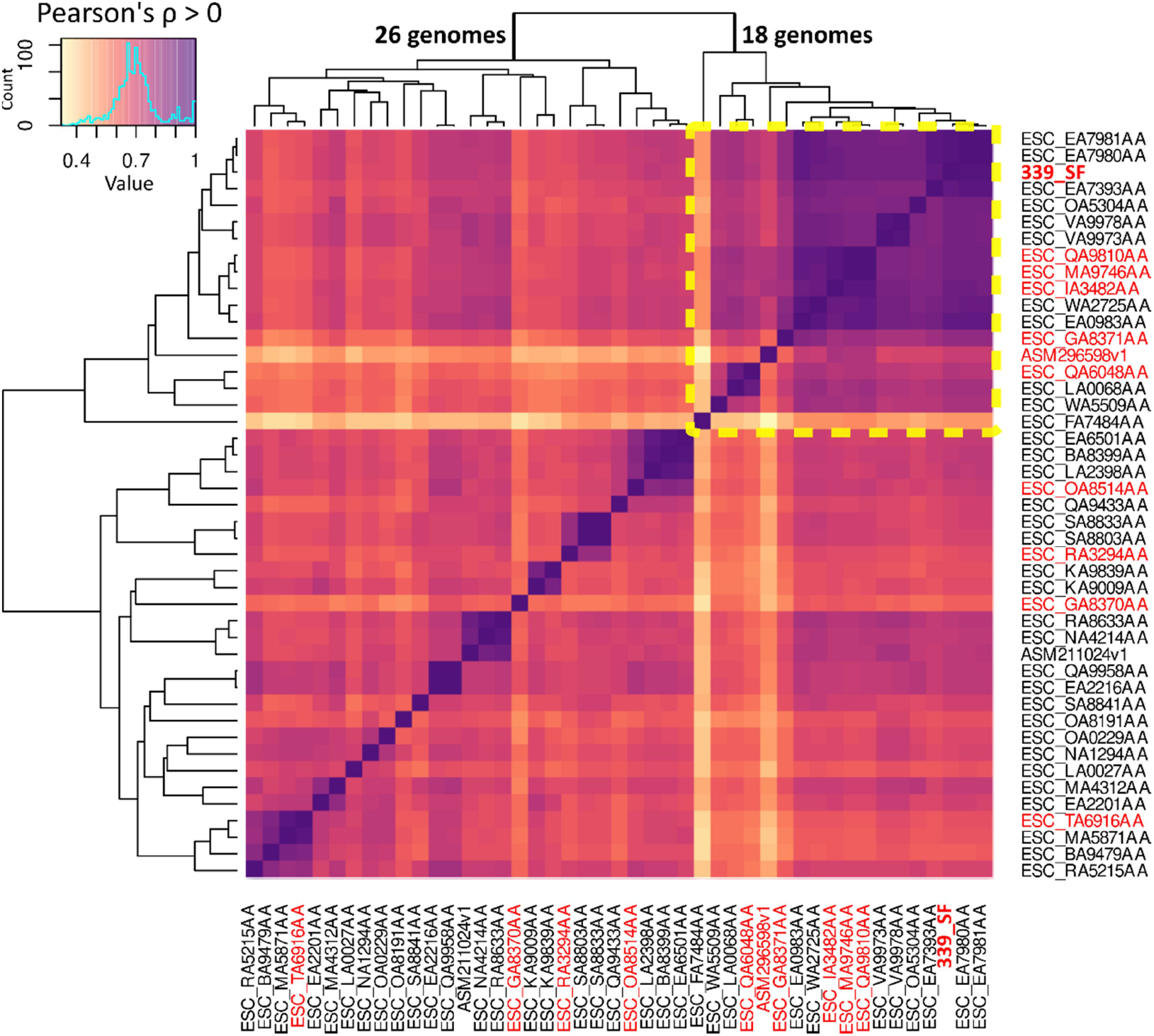
Hierarchical clustering of cryptic clade IV genomes based on functional genes presence/absence. The colours of the heat map indicate the Pearson correlation coefficient between 0 to 1 among the genomes. The colour of the label indicates the ecotype: red for environmental origin genomes and black for host origin. The cluster containing the 339_SF, labelled in bold red, is boxed in a yellow dotted line.

In parallel, we looked for genes enriched in clade IV genomes in relation to their ecotype (environmental or host). We found 24 over-represented genes in environmental genomes and only 3 over-represented genes in host genomes (**Table S3**). Of the 24 genes enriched in environmental ecotypes, 12 were previously described for *E. coli* reference strains, while the remaining 12 were identified in reference genomes belonging to other bacterial species. These genes may be linked to different key biological functions related to a free-living style. A group of factors are directly associated with the response to stress conditions and DNA damage: *UmuC* and *UmuD* help repair DNA damage and participate in the SOS response, *yrecE* and *ybcO* are nucleases that intervene in DNA repair, *dnaK* has chaperone activity during stress events and *kilR* encodes an inhibitor of cell division in response to antibiotics. The gene *hicA*, which encodes for a component of a type II toxin-antitoxin system in *E. coli*, has also been associated with the stress response. In addition, genes related to viral infections were also enriched in the environmental set: *cas6f* and *csy3* belong to the CRISPR defence system, *hpaIIM* encodes a restriction enzyme described in *Haemophilus parainfluenzae*, and *intQ*, is related to the integration of phages into bacterial genomes. Other enriched genes are the *bepC* genes, which confer antibiotic resistance to substances of hydrophobic nature, and *icsA*, which encodes a protein essential for bacterial adhesion and virulence (described in *Shigella flexneri*). Also, a group of genes associated with the environmental ecotype is linked to structural and transmembrane transport functions (*yidK, csbX, ompC, yidI, yknY, fliC* and *gpFI*). Finally, genes linked to energetic metabolism were also identified (*ahr*, which is involved in lipid metabolism, and *hypBA1*, related to carbohydrate metabolism).

On the other hand, over-represented genes among the host-origin genomes were *rrrD*, which encodes a factor with hydrolytic activity of bacterial walls, and *tolA*, which encodes a Tol-Pal system factor carrying key functions in cell division and outer membrane integrity. Both genes were previously described in *E. coli* reference strains. A third gene associated with host ecotypes is *prfA*, which was described in *Mycobacterium tuberculosis* as involved in protein biosynthesis.

## Discussion

Phylogenetic analyses combined with comparative genomics approaches can help clarify the evolutionary origins of environmental ecotypes. By conducting a comparative analysis of genomes within the *Escherichia* cryptic clade IV from diverse ecological (environmental or animal host) and geographical sources, we identified substantial intra-group genomic structuring linked to ecological origins. Our results shed light on the genomic mechanisms underlying the naturalization phenomena within the *Escherichia* genus and include the first genomic characterization of a member of *Escherichia* cryptic clade IV isolated from a highly polluted urban stream in South America. Furthermore, this study provides relevant data for the sanitary management of urban basins.

There is an ongoing debate about the ecological niches occupied by the cryptic clades and their phylogenetic relationships within the genus *Escherichia* (Vital *et al*., 2015; Walk, 2015; Jang *et al*., 2017). In this context, the cryptic clade IV genome reported here represents a particularly useful new data point, because it is the first isolate of this type obtained from an environmental source in South America, thus expanding the geographical representation of isolates from cryptic clades. Our analysis highlighted consistent relationships between cryptic clades and *E. coli* strains, which is coherent with previous phylogenies based on single nucleotide polymorphisms (SNPs) or other sets of genes (Walk *et al*., 2009; Walk, 2015; Jang *et al*., 2017; van der Putten and Mende, 2019). In addition, we observed that strains from clades III and IV fall in close phylogenetic proximity. It has been proposed that the latter groups comprise a new species, which was named *E. ruysiae* (van der Putten and Mende, 2019). Clade I strains clustered together with *E. coli* strains, many of which are known to be human pathogens. This observation aligns with previous studies and observational data, suggesting that clade I may represent a versatile taxon with intermediate properties between a free-living lifestyle and enteric commensalism (Chaudhuri and Henderson, 2012; Clermont *et al*., 2013).

Regarding genomic features of clinical and epidemiological interest, the genome of 339_SF and other clade IV genomes exhibited low abundance of virulence factors and no antibiotic-resistance genes were detected in most of the genomes of the clade. These characteristics, consistent with the findings of other authors on this clade (Ingle *et al*., 2011), have been associated with a free-living lifestyle (Casadevall and Pirofski, 2000; Uddin *et al*., 2021), as the distribution of these factors is typically linked to specific selection pressures associated with a host-associated life (Jernberg *et al*., 2010; Becattini *et al*., 2016).

Natural environments, such as urban freshwaters, present specific ecological conditions that act as filters for bacterial taxa. Conditions affecting the viability of microorganisms include a wide range of physicochemical characteristics (e.g., temperature, pH and nutrient availability) and stressors (e.g., interspecific competence, solar radiation and xenobiotic compounds) (Vrede, 2005; Jiang and Patel, 2008; Jang *et al*., 2017). Therefore, the success of bacterial taxa in persisting and proliferating in these environments depends on their physiological aptitude to face these ecological contexts. To identify genomic factors related to this differential fitness, we explored the enrichment of genes within the cryptic clade IV in relation to their origin (environmental or host). This approach allowed the identification of numerous genes likely associated with the free-living ecotype. Many of these genes function in stress response and DNA repair mechanisms. As well, elements associated with the CRISPR defence system were enriched in environmental genomes. These genes likely confer a selective advantage related to the reduction of high viral loads of open natural systems (Jackson *et al*., 2017). Additionally, genes linked to lipid and carbohydrate metabolic pathways were enriched in the environmental ecotype, suggesting that the utilization of alternative energy sources may be a feature of environmental isolates. Similar deductions were made by Luo et al. (2011), who also found enrichment of factors related to resource acquisition (diol utilization) and survival (lysozyme production; associated with bacterial innate immunity) in environmental strains of *E. coli*. Overall, the abundance of factors with defined actions against environmental stressors or adaptation to distinctive physicochemical characteristics is consistent with free-living ecotypes. Certainly, the limited range of ecological conditions expected in the enteric cavities of homeotherms would not select for such a wide array of genetic features.

We did not find differences in genomic features of clinical relevance among clade IV genomes in relation to their origin. In the same vein, neither the phylogenetic analysis nor the gene content-based clustering method completely discriminated clade IV genomes according to their origin. However, our results demonstrate that there are differences in gene content associated with the origin of isolates. These differences could be interpreted as evolutionarily selected traits, reinforcing the idea of ecological niche specialization. Thus, our results highlight a rich genomic diversity within the cryptic clade IV of the *Escherichia* genus. We believe that future work should be oriented to analyze the association of genomic features with phenotypic characteristics (e.g. growth, adhesion) on *Escherichia* isolates from different sources. This kind of studies will further clarify the relationship between genomic features and ecological niches in the *Escherichia* genus.

## Authors contributions

**Martín Saraceno:** conceptualization (equal contribution); formal analysis (head); investigation; writing – original draft preparation (head); writing – review & editing (head). **Nicolás Frankel:** formal analysis (supporting); project administration supervision (supporting); writing – review & editing (supporting). **Martín Graziano:** conceptualization (equal contribution); formal analysis (supporting); project administration supervision (head); writing – review & editing (supporting).

## Acknowledgements and conflict of interest statement

This work was financed by grants from CONICET (PUE 22920160100122CO). M.S. is recipient of a postdoctoral fellowship from CONICET. We thank Dr. David Kachanovsky and Dr. Nicolás Nahuel Moreyra for their assistance in phylogenomic analyses. The authors declare no conflict of interest.

## Data availability statement

The genome assembly for *Escherichia* cryptic clade IV strain 339_SF has been deposited at DDBJ/ENA/GenBank under the accession JBBUKW000000000 (BioProject PRJNA1092102, BioSample SAMN40619928). Accession information and metadata for the genomes of cryptic clade IV and the rest of the strains used are listed in Table S1.

